# Bioprinted pluripotent stem cell-derived kidney organoids provide opportunities for high content screening

**DOI:** 10.1101/505396

**Authors:** J. William Higgins, Alison Chambon, Kristina Bishard, Anke Hartung, Derek Arndt, Jamie Brugnano, Pei Xuan Er, Kynan T. Lawlor, Jessica M. Vanslambrouck, Sean Wilson, Alexander N. Combes, Sara E. Howden, Ker Sin Tan, Santhosh V. Kumar, Lorna J. Hale, Benjamin Shepherd, Stephen Pentoney, Sharon C. Presnell, Alice E. Chen, Melissa H. Little

## Abstract

Recent advances in the directed differentiation of human pluripotent stem cells to kidney brings with it the prospect of drug screening and disease modelling using patient-derived stem cell lines. Development of such an approach for high content screening will require substantial quality control and improvements in throughput. Here we demonstrate the use of the NovoGen MMX 3D bioprinter for the generation of highly reproducible kidney organoids from as few as 4,000 cells. Histological and immunohistochemical analyses confirmed the presence of renal epithelium, glomeruli, stroma and endothelium, while single cell RNAseq revealed equivalence to the cell clusters present within previously described organoids. The process is highly reproducible, rapid and transferable between cell lines, including genetically engineered reporter lines. We also demonstrate the capacity to bioprint organoids in a 96-well format and screen for response to doxorubicin toxicity as a proof of concept for high content compound screening.

## Introduction

The kidney plays a crucial role in the elimination of xenobiotics from circulation, regulation of blood pressure, and the maintenance of fluid, glucose, and electrolyte homeostasis ^1^. The combination of being highly perfused and the expression of a repertoire of specific transporters leads to increased xenobiotic exposure and risk of organ toxicity. As such, evaluation of nephrotoxicity is a key area of interest in drug development. Current preclinical screening tools for nephrotoxic compounds consist primarily of panels of human and animal renal proximal tubule epithelial cells (RPTEC) or small animal models. However, these systems often fail to accurately predict organ-specific toxicity, either as result of species-specific differences, or the inability to recapitulate relevant aspects of kidney physiology ^2^. While freshly isolated primary human RPTECs address the challenges of species specificity, the cells rapidly dedifferentiate in 2D and senesce when cultured in isolation, losing expression of key functional components including transporters and metabolic enzymes ^3-5^. To enable the effective development of drugs that are safe and effective for treating kidney disease, there is a critical need for modelling human kidney diseases and injury *in vitro* during preclinical drug development.

Human pluripotent stem cells (hPSCs) are a unique source of cells with which to model early development of mammals and the tissues resulting from directed differentiation represent a promising tool for drug development and therapeutic applications for human diseases. The unique nature of these cells lies in their capacity for unlimited self–renewal, organ-specific differentiation, receptivity to genetic alteration, and ease of access to phenotypic patient backgrounds through the process of reprogramming ^6^. Recent advances in the directed differentiation of hPSCs to human kidney cell types ^7-9^, including our own protocol for the formation of 3D kidney organoids containing all key renal progenitor lineages ^10-12^, raises the prospect of a single starting cell population. With scale up, such a differentiation protocol could enable high content drug screening in the near term, and the generation of functional kidneys for transplantation in the long term.

Tissue engineering using 3D bioprinting enables the fabrication of tissues through an automated, spatially controlled, and reproducible deposition of living cells in defined geometric patterns ^13-16^. In addition to establishing some key features of native tissue architecture *in* vitro, bioprinting may also enhance reproducibility and provide finer resolution and smaller scale than is achievable manually. Here we demonstrate the successful adaptation of our directed differentiation protocol using the NovoGen MMX bioprinter to achieve: (1) automated fabrication of self-organizing kidney organoids equivalent at the level of morphology, component cell types and gene expression to those previously reported via manual generation; (2) the generation of large numbers of uniform and highly reproducible organoids in reduced time; (3) the generation of bioprinted organoids from distinct hPSC cell lines, including gene-edited reporter lines, (4) adaptation of fabrication protocols to a 96-well plate format amenable to compound efficacy and safety screening and (5) proof-of-concept for the application of bioprinted organoids for toxicity screening.

## Results

### Automated fabrication of kidney organoids via bioprinting

The directed differentiation protocol being adapted for biofabrication has been previously described ^10-12^. In brief, hPSCs cultured in a monolayer format are initially induced to commit to posterior primitive streak via the addition of the potent GSK3β inhibitor, CHIR99021 to activate canonical Wnt signalling, for 4 days then transitioned to intermediate mesoderm via the addition of FGF9 for a further 3 days. At this time, all cells are enzymatically dissociated, centrifuged to facilitate reaggregation and then manually transferred to Transwell^®^ permeable supports for culture as a micromass at an air-media interface. The objective of this study was to reproduce the original protocol from the completion of monolayer culture at Day 7, thereafter replacing the manual process of micromass formation with bioprinting. After the initial steps of 2D differentiation (7 days) and cellular dissociation, cells were either centrifuged to generate cell pellets for manual transfer to Transwell permeable supports or transferred as a cell only paste into syringes in preparation for bioprinting. Initial bioprints were performed with an in-syringe centrifugation of cellular material (“bioink”) for compaction and the removal of any trace media (dry ink) (Figure 1A). Upon maturation, all dry ink organoids exhibited large regions of non-nephric differentiation and lacked higher level 3-dimensional organization, regardless of whether they were printed to match the diameter of manually prepared organoids or to target a specific cell number in size (Figure 1A). To assess whether differentiation outcome could be improved by the addition of small volumes of media to ease mechanical stress on the cells and to facilitate uniform deposition, the in-syringe centrifugation step was omitted and the bioink was diluted slightly by the addition of media with gentle mixing using a pipette prior syringe loading. Histological evaluation of bioprinted organoids generated with wet bioink closely resembled those derived from manual preparations. Nephron differentiation patterns in manual and bioprinted organoids were comparable by brightfield microscopy and after histological sectioning, using either Hematoxylin & Eosin (H&E) staining or immunofluorescent staining for markers such as LTL^+^/ECAD^+^ (Figure 1A). Organoids derived from dry bioink were largely devoid of the glomerular and tubular structures observed within organoids either manually generated or printed using wet bioink. Histological analysis of sections through bioprinted organoids revealed a stromal bed along the Transwell above which a network of interconnected nephrons was evident (Figure 1B). Immunofluorescent staining identified MAFB+ podocytes in putative glomeruli, LTL^+^ proximal tubules, ECAD^+^ tubule structures, and a contiguous GATA3^+^/ECAD^+^ collecting duct (Figure 1BCDE). The spatial arrangement of these kidney markers aligned closely with previously reported manually produced kidney organoids ^10-12^, with glomerular markers in the upper regions of the organoid and transition from proximal tubule to collecting duct towards the bottom of the organoid (Figure 1BCDE).

**Figure 1.**
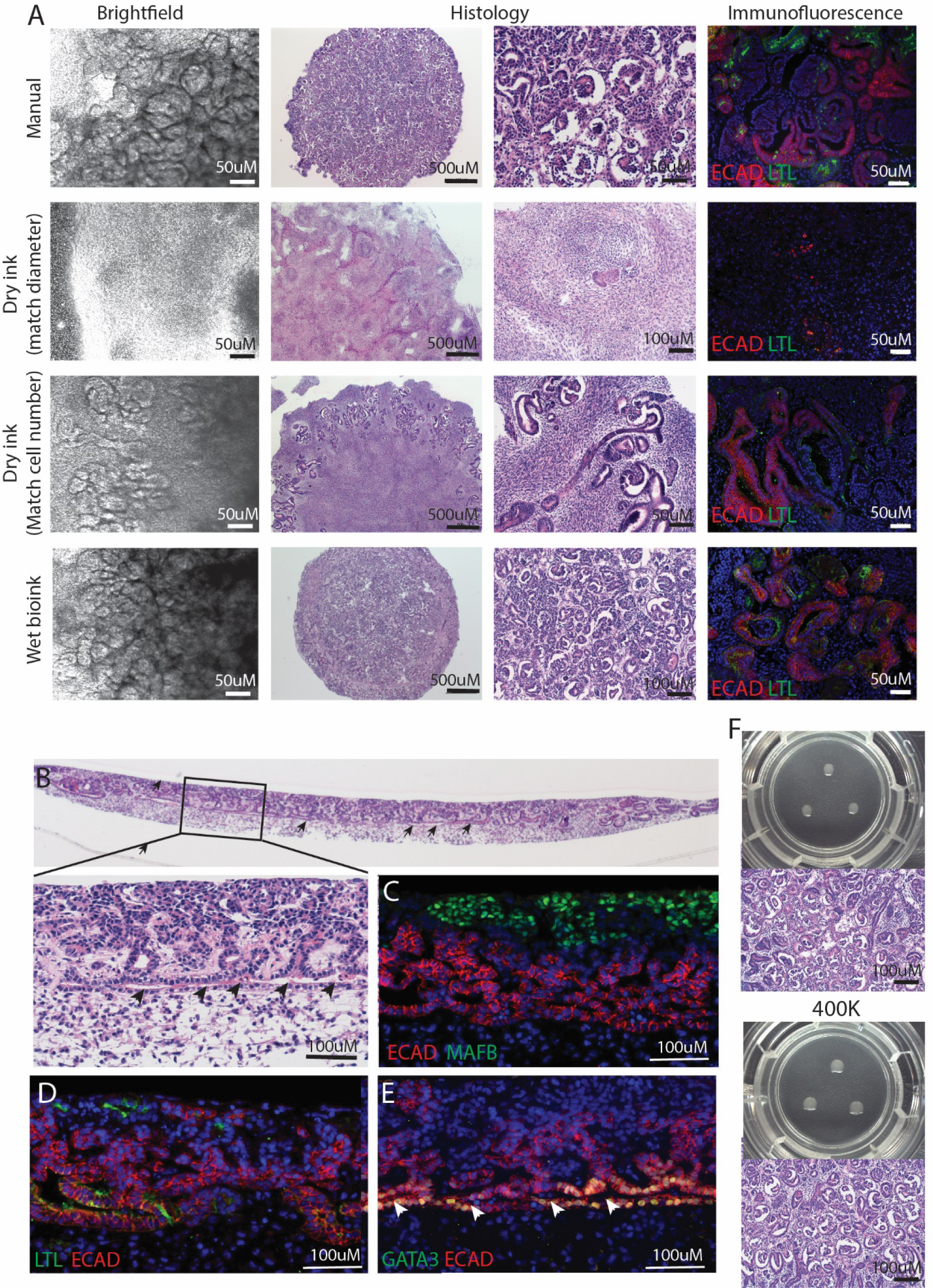
Optimisation of bioprinting methodology for the generation of kidney organoids. **A.** Brightfield, histological and immunofluorescence comparisons of kidney organoids generated manually (5 × 10^5^ cells per organoid), using dry cell paste (ink) controlled for organoid diameter, dry ink controlled for cell number and wet bioink. **B.** Histological cross-section of bioprinted organoids showed a high level of tissue organisation. **C.** MAFB+ podocytes reside in close proximity to ECAD+ tubular regions. **D.** LTL+ECAD+ proximal tubular regions are present. **E.** A contiguous putative collecting duct (GATA3+/ECAD+) network spans horizontally throughout organoids with nephrons connected to and contiguous with this epithelium. **F.** Kidney organoid differentiation within bioprinted organoids is equivalent even with reduced starting cell number. Images show three kidney organoids bioprinted within a single Transwell^TM^, together with the resulting histology, from either 2 × 10^5^ or 4 × 10^5^ starting cells harvested from a Day 7 differentiation culture.

### Bioprinting provides improved reproducibility, increased throughput and higher organoid yield

To evaluate the effect of cell concentration, organoids were printed from cell pastes ranging in concentration from 200,000 cells/µL to 400,000 cells/µL (Figure 1F) with both cell densities resulting in efficiently differentiated organoids. Indeed, appropriately patterned organoids could be generated using a considerable range of total printed cell numbers with the bioprinter being programmed to deposit between 1 and 9 organoids per Transwell in a 6 well plate (Figure 1F; Suppl. Fig. 1B). The NovoGen bioprinter reproducibly printed organoids ranging in calculated cell number from 50,000 to 500,000 cells per organoid simply by adjusting the volume of bioink extruded (Suppl. Fig. 1A). The smallest organoid printed was estimated at 4000 cells based upon loading cell density (100,000cells/µl) and volume (0.04µl) (Suppl. Fig 1C), however reproducibility at this volume and cell density was lower.

NovoGen Bioink cell density ranged from as low as 10,000 cells/µl up to 400,000 cells/µl. Optimal organoids were generated via a displacement of 0.48µl of a 200,000 cells/µl bioink (approximately 96,000 cells per organoid at time of print). Organoids of reduced size maintained similar differentiation capacity as compared to larger sized organoids (Suppl. Figure. 1C).

Takasato et al ^11^ described a manual protocol for kidney organoid generation using 500,000 cells. Reductions of total cell numbers, while possible, was limited by the manual dexterity required to both see and handle smaller pelleted organoids. As automated fabrication enabled software controlled volumetric dispensing, the reproducibility of kidney organoids generated using the NovoGen bioprinter was evaluated across a range of total cell numbers. Post-print images were captured and the diameters of the printed organoids were measured, demonstrating minimal variability in size (coefficient of variation; % CV) across each size group (Table 2).

**Table 1.** Details of antibodies and lectins used for immunofluorescence of organoids.

| **Specificity** | **Host species** | **Manufacturer and identifier** | **Dilution range** |
| --- | --- | --- | --- |
| **MAFB** | Mouse | Novus Biologics (NBP2-45718) | 1:400 |
| **NEPHRIN** | Sheep | R&D Systems/Bioscientific (AF4269) | 1:100-1:300 |
| **E-CADHERIN** | Mouse | Abcam (ab1416) | 1:100 |
|  | Mouse | BD Biosciences (610181) | 1:300 |
|  | Rabbit | Abcam, ab15148 | 1:100 |
| **GATA3** | Mouse | Invitrogen (MA1-028) | 1:100 |
|  | Goat | R & D Systems (AF2605) | 1:300 |
| **CD13**<br>**(PerCP-Cy4.5 conjugate)** | Mouse | Australian Biosearch (301713) | 1:300 |
| **CUBILIN** | Goat | Santa Cruz Biotechnology (sc-20607) | 1:300 |
| **EpCAM**<br>**(Alexa488 conjugate)** | Mouse | Biolegend (324210) | 1:300 |
| **LAMININ** | Rabbit | Sigma-Aldrich (L9393) | 1:400-1:500 |
|  | Rabbit | Abcam (ab11575) |  |
| **MEIS1/2/3** | Mouse | Australian Biosearch (ATM39795) | 1:300 |
| **Proximal tubule brush border membrane** | Lotus tetragonobulus lectin (LTL) | Vector Laboratories (B-1325) | 1:300-1:500 |
| **SLC12A1** | Rabbit | Proteintech (18970-1-AP) | 1:300 |
| **CLEAVED CASPASE 3** | Rabbit | Cell Signaling Technology (9661) | 1:400 |
| **CYTOKERATIN 8/18** | Guinea Pig | Abcam (ab194130) | 1:400 |

**Table 2.** Reproducibility of Deposition.

| **Organoid Size** | **Volume Bioprinted** | **Organoid Number** | **Mean Diameter of Deposit (mm)** | **%CV** |
| --- | --- | --- | --- | --- |
| 100K | 0.49 μL | 24 | 1.79 | 3.68 |
| 200K | 0.98 μL | 24 | 2.30 | 1.08 |
| 500K | 2.43 μL | 24 | 3.12 | 2.93 |

One advantage of bioprinting is the speed with which tissues can be generated. Manual processing allows for the generation of approximately 30 organoids per hour and only a maximum of 3 organoids can be placed on a filter. The efficiency with which kidney organoids can be generated using the NovoGen bioprinter was evaluated by timing each aspect of the process. The total time required to bioprint 108 organoids was 10 min, 38 seconds, a time 15-20 times faster than manual production of the same number and configuration of organoids. Taken together, the NovoGen bioprinter not only automated a manual process, but also increased control of organoid size, allowing for a greater number of organoids to be printed from the same starting material, enabling increased throughput, uniformity, and additional endpoint evaluation.

### Bioprinted organoids show complex morphology consistent with a model of developing human kidney

Further characterisation of kidney differentiation within bioprinted organoids was performed using wholemount brightfield, immunofluorescence and live fluorescence imaging. After printing, brightfield examination of bioprinted organoids revealed the formation of tubular structures by Day 7+ 5 (7 days of monolayer culture + 5 days organoid culture post bioprinting) with increasing complexity from thereon (Figure 2A). Simultaneous four colour staining showed the presence of contiguous nephron epithelia surrounded by a LAMININ^+^ basement membrane and segmentation into glomeruli (NEPHRIN), proximal segment (LTL, CUBN, CD13), distal segment (ECAD, EPCAM, SLC12A1) and collecting duct (GATA3, EPCAM, ECAD) (Figure 2BC). The nephrons were surrounded by a MEIS1/2/3 positive stroma and an intervening endothelium (CD31) (Figure 2BC).

**Figure 2.**
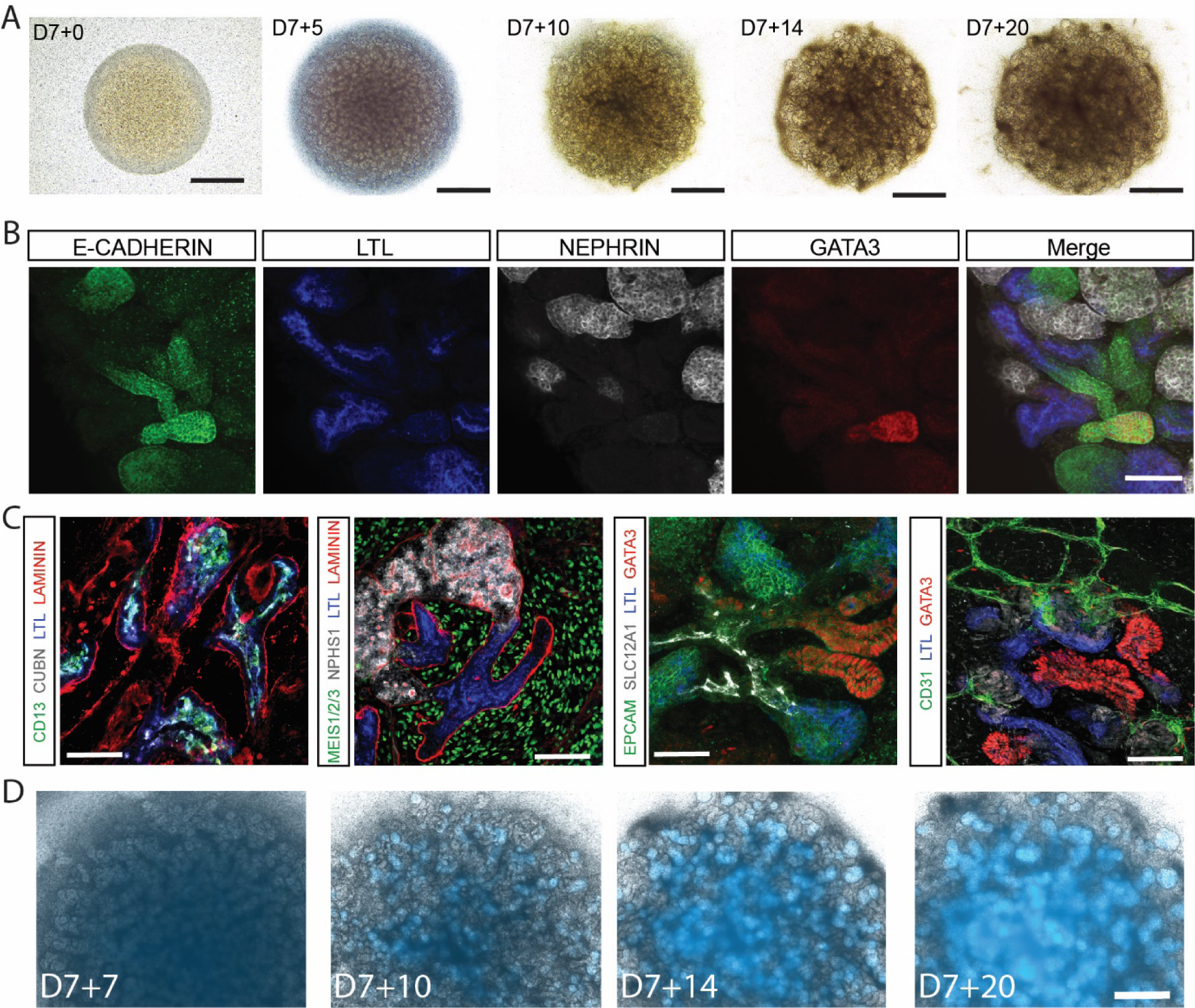
Characterisation of kidney organoids bioprinted using control and reporter iPSC lines. **A.** Brightfield images of bioprinted organoids across time showing evidence of increasing tubular complexity. Scale bar represents 800 µm. **B.** Immunofluorescence of a Day 7+18 bioprinted organoid showing the presence of nephron epithelium (E-CADHERIN, green), proximal tubules (LTL, blue), collecting duct (GATA3, red) and podocytes (NEPHRIN, grey). Scale bar represents 100µm. **C.** Immunofluorescence of Day 7+18 bioprinted organoids stained for the proximal tubule markers CD13, LTL and CUBN, podocyte marker NPHS1, distal tubule epithelial markers EPCAM and SLC12A1, collecting duct markers EPCAM and GATA3, the presence of a LAMININ-positive basement membrane along the nephrons, a surrounding MEIS1/2/3-positive stroma and CD31-positive endothelium. Scale bar represents 50µm. **D.** Bioprinted organoids generated using a MAFBmTAGBFP iPSC reporter line showing the formation of glomeruli containing MAFB+ (blue) podocytes across time. Scale bar represents 200µm.

The capacity to generate large numbers of kidney organoids using automated fabrication provided an opportunity for high content screening with a rapid image readout. To evaluate how effectively this can be performed, organoids were bioprinted using CRISPR/Cas9 gene-edited reporter iPSCs. The MAFB^mTagBFP2^ reporter line has been previously used to monitor commitment to a podocyte lineage, allowing the visualisation of glomeruli in live organoids *in vitro* ^17^ and after organoid transplantation ^18^. Organoids were bioprinted onto Transwells (1 × 10^5^ cells per organoid, 9 organoids per well) and flourescent microscopy was used to monitor the onset and upregulation of blue fluorescent protein (BFP) expression from the endogenous MAFB locus at multiple timepoints during organoid development (Figure 2D; Suppl Figure 2). BFP-expressing podocytes were observed approximately 10 days post organoid formation (d7+10) and persisted thereafter (Figure 2D). The same reporter line was then used to evaluate the effect of cell number on differentiation. Organoids were bioprinted using final cell numbers of 4 × 10^3^ and 5 × 10^4^ and 1 × 10^5^ cells per micromass and imaged live at two timepoints across differentiation (Suppl. Figure 2). The identity of tubular epithelium was uniform irrespective of the number of cells used for bioprinting (Suppl. Figure 2). The use of such a reporter line obviates the need for antibody staining and demonstrates that live fluorescence imaging of bioprinted organoids is feasible.

### Equivalence of cellular composition and transcriptional identity between standard and bioprinted organoids

While histological and immunofluorescence data suggested the presence of equivalent cellular components in organoids generated via bioprinter fabrication, this was examined in more detail by directly comparing the transcriptional profile of single cells isolated from a standard organoid versus a bioprinted organoid, both generated using the same iPSC line, CRL1502.C32. The standard organoid was generated using 5 × 10^5^ cells while the bioprinted organoid was generated with only 4 × 10^3^ cells (Suppl. Figure 1B). Single cells were captured using the 10x Chromium platform and sequenced using an Illumina HiSeq. Data from both samples was combined and clustered in Seurat v2.3.1 ^19^ to generate a single aligned dataset to facilitate a direct comparison between the two starting organoids (Figure 3A). Seven distinct clusters were identified, including Cluster 4, in which cells showed differential expression of nephron markers including PAX2, PAX8 and MAFB, and Cluster 5 which showed differential expression of endothelial markers, including PECAM1 and SOX17 (Figure 3ABCD; Suppl. Figure 3ABC; Suppl. Table 1). Across all clusters, there were similar proportions of cells from each organoid represented, indicating strong equivalence between manual and bioprinted organoids and between organoids generated with large or small numbers of starting cells (Figure 3E). Reclustering of Cluster 4 allowed the identification of committing nephron progenitor (DAPL1, LYPD1, SIX1), early tubule (LHX1, HNF1B, EMX2), and podocyte (MAFB, PODXL, TCF21) cell types (Figure 3F; Suppl. Figure 3DE; Suppl. Table 2) with comparable proportions of cells from each organoid present in each of these subclusters (Figure 3D) and equivalent transcriptional identity between cells from the standard and bioprinted organoids for each subcluster (Figure 3G; Suppl. Figure 3E; Suppl. Table 2). This again shows that bioprinted organoids are equivalent to standard organoids at the level of cellular complexity and identity.

**Figure 3.**
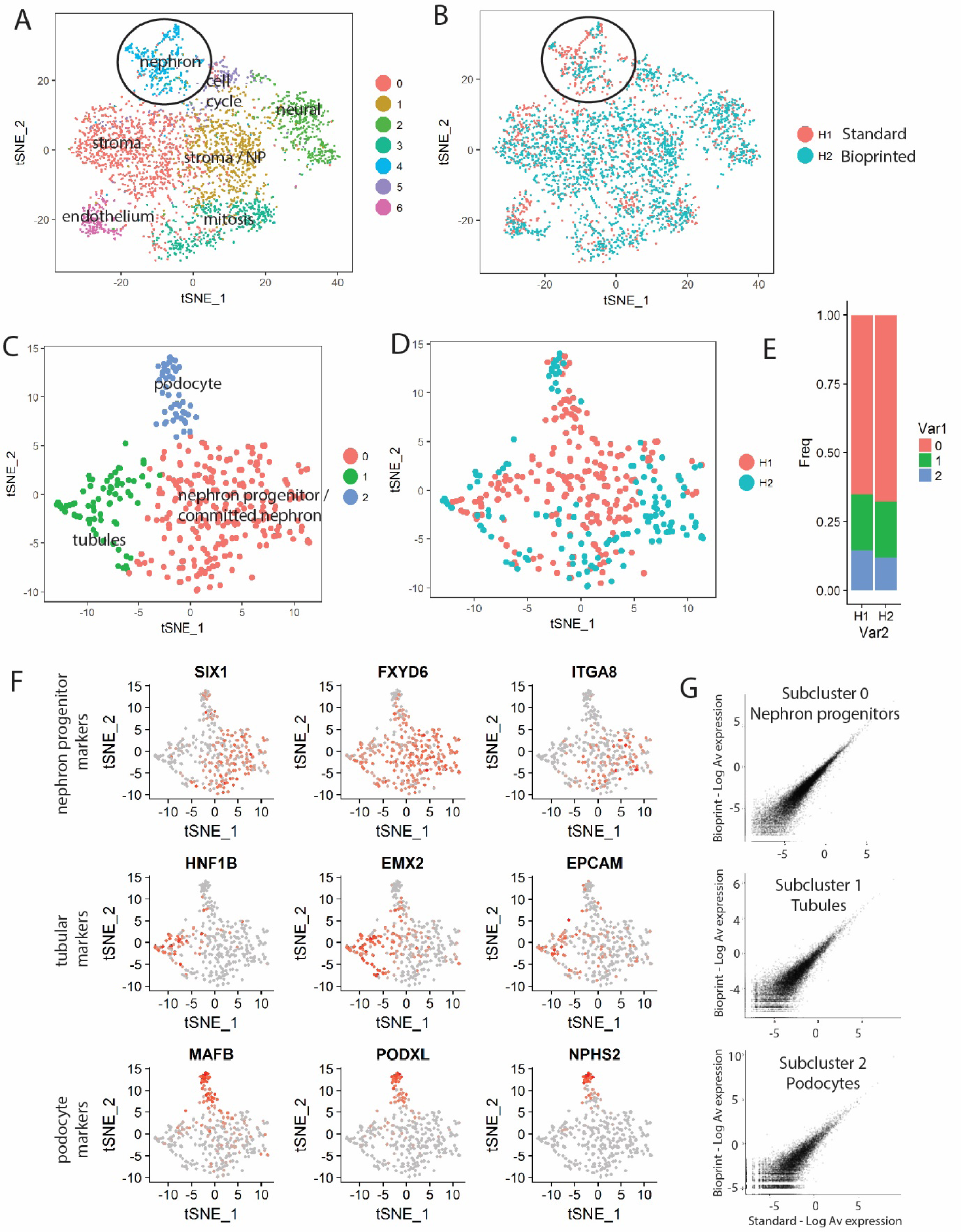
Single cell transcriptional profiling shows equivalence between standard and bioprinted kidney organoids. **A.** tSNE overlay of 3671 cells isolated from manual 1290 cells) or bioprinted (2381 cells) kidney organoid cells. Unsupervised clustering with Seurat identified 7 distinct cell clusters and direct comparison with standard organoids, identified as indicated using GO analysis and comparison to available human fetal kidney data ^29^. **B.** A view of the same tSNE plot indicating whether individual cells were derived from the manual (pink) or bioprinted (blue) organoids. **C.** Reanalysis of the nephron cluster (cluster 4) reveals the presence of three subclusters, identifiable as committed progenitor / renal vesicle (subcluster 0), early tubule (subcluster 1) and podocyte (subcluster 2). **D**. tSNE plot identifying nephron cluste-+r cells based upon their origin in either manual (pink) or bioprinted (blue) organoids. **E.** Relative ratios of cells present in each subcluster for manual and bioprinted organoids. **F**. Feature plot showing cells expressing key differentially expressed genes for each of the nephron progenitors (Subcluster 0; SIX1, FXYD6, ITGA8), tubular epithelium (Subcluster 2; HNF1B, EMX2, EPCAM) and podocytes (Subcluster 2; MAFB, PODXL, NPHS2). Expression indicated by colour: low (grey)-high (red).**G.** A comparison of average gene expression values (each point is a gene) within each nephron subcluster between manual and bioprinted organoids shows tight transcriptional congruence.

### Application of bioprinted organoids in toxicity screening

Doxorubicin, an anthracycline antibiotic used for cancer treatment, has been shown to induce glomerular nephrotoxicity both *in vitro* and clinically ^20-23^. To assess the capacity of 3D bioprinted kidney organoids to model drug-induced glomerular toxicity, organoids were treated with 2µM or 10 µM of doxorubicin for 72 hours (Figure 4A). Histological assessment by H&E staining of untreated control organoids revealed the presence of distinct glomerular and tubule structures as described above, whereas Dox-treated organoids exhibited concentration-dependent morphological changes consistent with cell injury and degeneration (Figure 4B). Organoids treated with low dose (2 µM) doxorubicin exhibited partial collapse of glomerular structures and tubule networks, whereas organoids treated with high dose (10 µM) doxorubicin exhibited near complete collapse of glomerular structures and tubule networks across the plane of section. Control organoids and organoids dosed with 10 µM doxorubicin were subsequently co-stained for Cleaved Caspase 3 (CC3) and either MAFB or LTL to identify regions undergoing apoptosis in response to treatment (Figure 4C). Control organoids displayed clear podocyte (MAFB^+^) and proximal tubule (LTL^+^) regions closely associated with pan-epithelial markers CK8+18, and were devoid of CC3 staining. Organoids treated with doxorubicin retained CK8/18 and LTL staining but lacked MAFB staining. Furthermore, the collapsed glomerular structures observed with H&E staining were found to be CC3^+^, suggestive of podocyte loss by apoptosis. Extended analysis by gene expression showed a dose-dependent increase in the apoptosis regulator, BCL2 Associated X protein (*BAX*), and increased caspase 3 (*CASP*) expression in response to doxorubicin treatment (Figure 4D). Kidney Injury Molecule-1 (*HAVCR*) was also upregulated in a dose-dependent manner, potentially due to the restructuring of the proximal tubule region around collapsed glomerular networks. Cell type-specific sensitivity to doxorubicin was further demonstrated by preferential downregulation of podocyte markers nephrin (*NPHS1*) and podocalyxin-like protein 1 (*PODXL*), compared to the proximal tubule marker, cubulin (*CUBN*). Together, these data provide evidence that iPSC-derived kidney organoids can display cell-type specific, drug-induced toxicity, supporting the utility of 3D bioprinted kidney organoids for use in glomerular toxicity screening.

**Figure 4.**
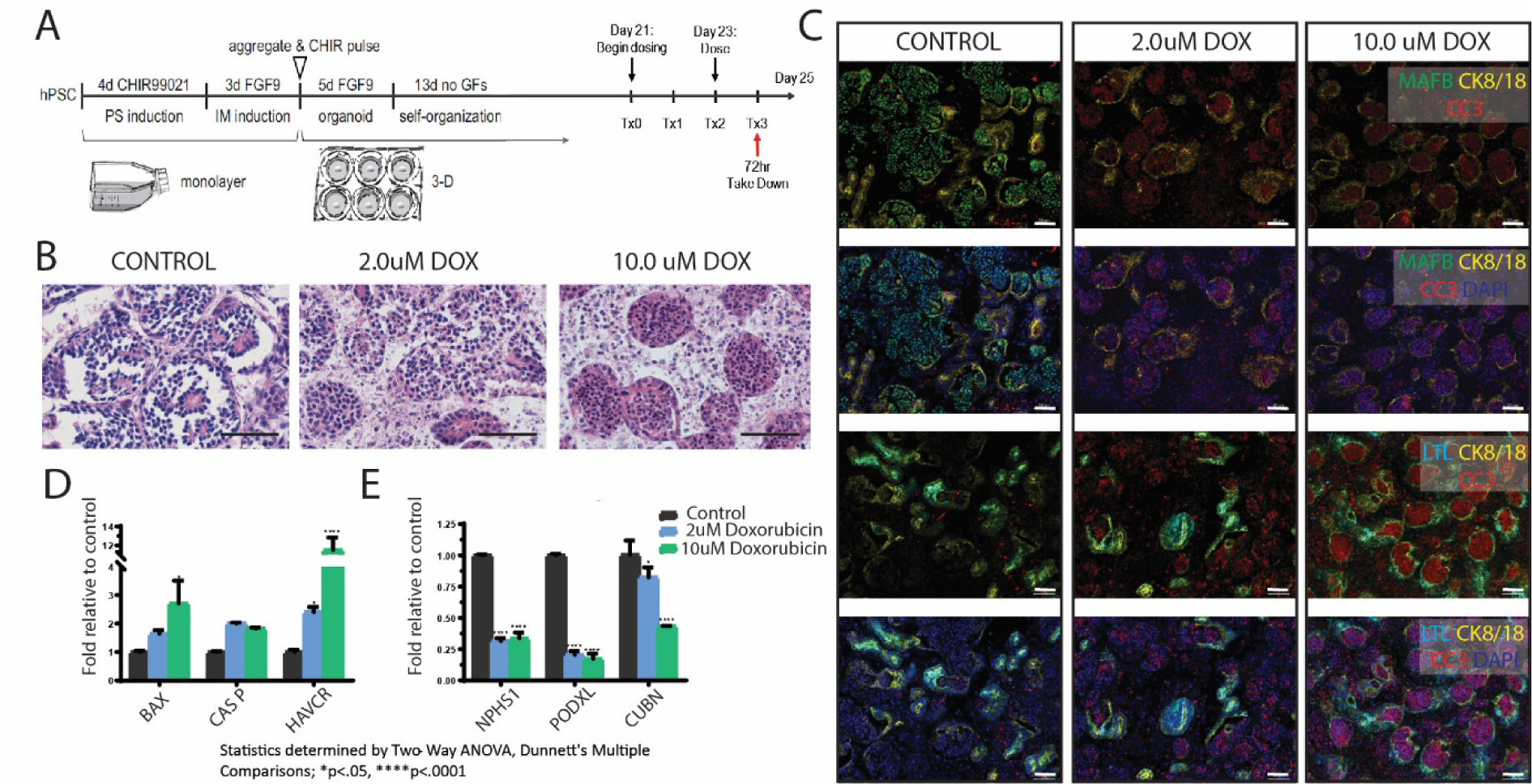
Application of bioprinted organoids in toxicity screening. **A.** Schematic outline of 2D differentiation, generation of 3D organoids by bioprinting, doxorubicin addition and analysis. **B.** H&E stains of sectioned organoids exposed to 72hr of 2.0 µM or 10.0µM doxorubicin treatment. **C.** Immunofluorescent staining of sectioned organoids after doxorubicin exposure for 72 hrs. Podocytes (MAFB, green); epithelium (CK8/18, yellow); proximal tubules (LTL, light blue), Apoptosis (CC3, red), DNA (DAPI, dark blue). **D.** Doxorubicin treatment leads to up-regulation of apoptosis and kidney injury genes (BAX, CASP, HAVCR). E. Preferential down-regulation of nephron markers (NPHS1, PDXL) compared to the proximal tubule marker (CUBN) upon doxorubicin treatment.

### Bioprinting kidney organoids in a 96-well plate format

To better utilize the growth area of the Transwell surface and increase downstream throughput for high content applications, we extended the methodology to 96-well printing of kidney organoids using a custom plate carrier (Figure 5A). As printing 96 organoids increased print time over the standard 18 organoids per 6-well plate, we evaluated cell viability after every two rows (or 24 organoids printed). Viability of the iPSC bioink remained high at 93-99% over multiple syringes (Figure 5B), thereby minimizing concerns about cell health with increased print time per plate. Maturation of kidney organoids cultured in 96-well Transwells displayed similar morphological progression over time to larger culture formats. The organoids printed as uniform circular micromasses of cell material across the whole 96-well plate (Figure 5CD) and showed evidence of differentiation at multiple time points that closely resemble manually prepared organoids at the whole tissue macroscopic level and by histological analysis via H&E (Figure 5C). All key nephron segments including glomerular podocytes (MAFB), proximal tubule cells (LTL, HNF4A) and distal tubules (ECAD), as well as putative collecting duct (GATA3/ECAD) were present (Figure 5E). To evaluate the application of bioprinted kidney organoids in 96-well format for drug toxicity screening, organoids were treated with a broad range of doxorubicin concentrations (0.2 to 25 µM) for 72 hours and harvested for viability assessment. Analysis of 6-well and 96-well viability results suggested that organoids printed in both plate formats were similarly sensitive to doxorubicin toxicity. The doxorubicin IC50 for organoids bioprinted in 6-well plates was 3.9 ± 1.8 µM (value ± standard error), while the calculated IC50 for organoids bioprinted in a 96-well plate was 3.1 ± 1.0 µM (Figure 5F). These results suggest that bioprinted kidney organoids cultured in 96-well Transwells closely recapitulate the morphology and differentiation capacity of both manually prepared organoids and organoids printed in larger plate sizes. In addition, the increased throughput afforded by utilizing a 96-well platform does not alter sensitivity to a doxorubicin, an established nephrotoxic drug.

**Figure 5.**
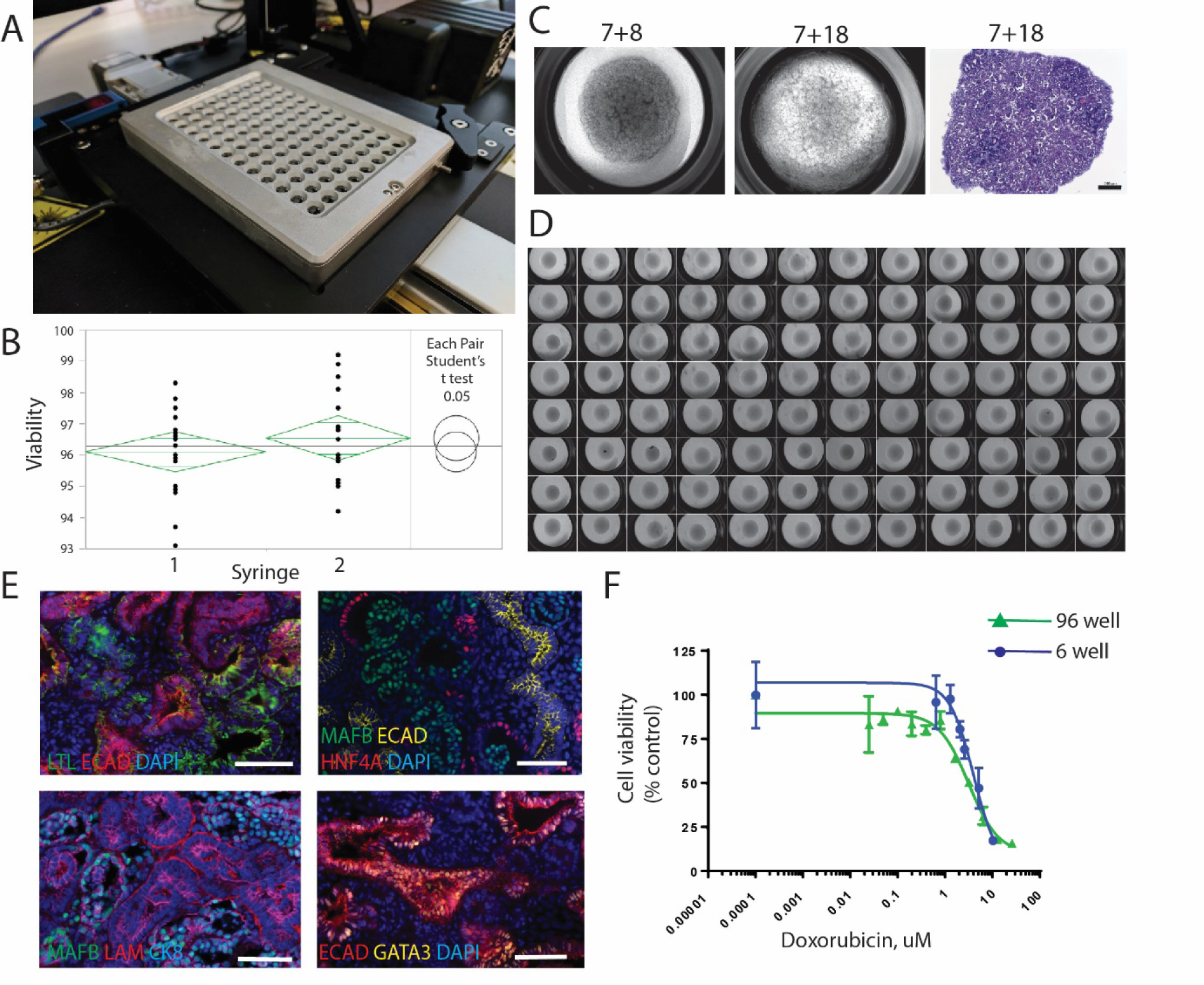
Bioprinting into 96-well plates for high content screening of compounds. **A.** Prototype 96-well anti-rotational plate to facilitate printing on Transwell permeable supports. Viability remains high (93-99%) across multiple rows of the plate and two independent bioink-containing syringes, demonstrating cell stability throughout the full 96-well bioprinting process. **C**. Brightfield and histology of kidney organoids bioprinted in 96-well format. Scale bar = 200uM. **D.** Kidney organoids printed on 96-well Transwell permeable supports. **E.** Immunofluorescent staining of bioprinted kidney organoid sections reveal the presence of proximal tubule (LTL), tubular epithelium (ECAD, HNF4), podocyte (MAFB), and putative collecting duct (ECAD/GATA3) marker expression. Scale bar = 50uM. **F.** Viability of cells within organoids plated in either 6-well or 96-well format in response to 72hr Doxorubicin.

### Discussion

This report describes an automated approach to kidney organoid generation via the use of 3D bioprinting. The kidney organoids generated using this mode of manufacture showed improved reproducibility among organoids, greater fidelity between independent starting cell lines, and equivalence to the gold standard manual protocol for generating kidney organoids at the level of cellular complexity, identity and gene expression. Importantly, the use of the bioprinter facilitated the generation of substantially larger numbers of organoids from the same number of starting cells by allowing the generation of organoids from as few as 4000 cells. Customized bioprinting scripts also allowed for the placement of 3 to 9 organoids per Transwell as well as the bioprinting of organoids in a 96-well format. Using fluorescent reporter iPSC lines designed to mark the identity of characteristic kidney cell types, we also show a proof of principle with respect to high content live imaging for maturation. As such, this suggests significant utility for bioprinted organoids in high content to medium throughput screening.

This is not the first report of the automated generation of human pluripotent stem cell derived kidney structures. Czerniecki et al ^24^ previously reported the use of a Tecan liquid handling platform for the distribution of undifferentiated hPSCs prior to differentiation and the automated image capture of nephron features across a 96-well format. The approach described here has several points of difference from that the previous study, including the protocol for directed differentiation and the increased histological complexity of the resulting tissue. The approach to bioprinting of kidney organoids described here generates a multicellular 3D structure containing nephrons, surrounding stroma and endothelium, as has been previously described using this differentiation protocol. Immunocytochemical evaluation of these bioprinted structures shows clear histological evidence of and interconnected epithelial network linking nephrons via a GATA3+ECAD+ tubular network. The cellular complexity and morphology was reproduced in bioprinted organoids generated using a variety of starting cell numbers. The equivalence of bioprinted organoids to those generated manually was reinforced at the level of single cell transcriptional profiling. Importantly, the use of the bioprinter facilitated the generation of large numbers of kidney organoids with a low %CV, as would be required for screening purposes. The bioprinting of 108 kidney organoids required approximately 10 minutes of time (648 per hour). By comparison, a skilled practitioner could not generate more than 30 organoids per hour manually. Importantly, the tissues generated can be processed for the analysis of complex histological readouts and the use of reporter iPSC lines to generate bioporinted organoids provides a fluorescence readout facilitating rapid and quantitative image capture. As a proof of concept for the use of this approach for high content compound screening, doxorubicin toxicity was assessed using organoids arrayed in either a 6 well or 96 well format. Hence, bioprinted organoids are likely to be useful for toxicity screening as well as the assessment of more complex disease phenotypes present when using patient-derived iPSC lines.

## Acknowledgements

Bioinformatic advice was provided by Alicia Oshlack and Luke Zappia, Murdoch Children’s Research Institute. We thank the Australian Genome Research Facility for access to 10x Chromium single cell library preparation and the Murdoch Children’s Research Institute Translational Genomics Unit for provision of Next Generation Sequencing. MHL is a Senior Principal Research Fellow of the National Health and Medical Research Council, Australia (APP1136085). This work was supported by Organovo Inc, California’s Stem Cell Agency grant number EDUC2-08388 and the NHMRC (GNT1100970, GNT1098654).

## Materials and Methods

### 2D iPSC Culture and Manual Organoid Production

All iPSC culture and differentiation procedures were performed with the iPS cell line CRL1502.C32, following published methodologies with slight modifications ^10-12^. In brief, human iPSCs were thawed and seeded overnight in the presence of 1x RevitaCell (ThermoFisher Scientific catalog# A2644501), and cultured under standard feeder-free, defined conditions on GelTrex (ThermoFisher Scientific catalog# A1413301) in Essential 8 medium (Thermo), with daily media changes. On the day prior to initiation of differentiation, the cells were dissociated with TrypLE Select (ThermoFisher Scientific catalog#12563011), counted using trypan exclusion on a Nexcellom Cellometer Brightfield Cell Counter (Nexcelom Biosciences), and seeded in a GelTrex coated T-25 flask in Essential 8 medium containing 1x RevitaCell (ThermoFisher catalog#A2644501). Intermediate mesoderm induction was performed by culturing iPSCs in STEMdiff APEL medium (STEMCELL Technologies catalog# 5210) or TeSR-E6 medium containing 6-8 μM CHIR99021 (R&D Systems catalog# 4423/10) for four days. On Day 4, cells were differentiated in STEMdiff APEL medium or or TeSR-E6 medium supplemented with 200 ng/mL FGF9 (R&D Systems catalog# 273-F9-025) and 1 μg/mL Heparin (Sigma Aldrich catalog# H4784-250MG). On Day 7, cells were dissociated with Trypsin EDTA (0.25%, Thermo Fisher catalog# 25200-072). The resulting suspension was counted with a Nexcelom Cellometer to determine the viable cells by trypan exclusion. Approximately 500,000 cells were then transferred into individual 1.5 mL sterile microcentrifuge tubes and centrifuged three times in a fixed rotor micro-centrifuge (180° rotation between spins) at 400 x *g* for 3 minutes. Resulting organoid pellets were transferred to 0.4 µm polyester membranes of 6-well Transwell permeable supports (Corning Costar catalog# 3450) with ART wide bore filtered pipette p200 or p1000 tips (Molecular Bioproducts catalog# 2069 and 2079, respectively). 3D organoids were cultured for 1 hour in the presence of 5 to 10 μM CHIR99021 in either STEMdiff™ APEL or TeSR-E6 medium in the basolateral compartment and subsequently cultured until Day 12 in STEMdiff™ APEL or TESR-E6 medium supplemented with 200 ng/mL FGF9 and 1 μg/mL Heparin (media only in the basolateral compartment). From Day 12 to Day 25, organoids were grown in STEMdiff™ APEL of TeSR-E6 media medium without supplementation. Kidney organoids were cultured until harvest at Day 25.

### Bioprinting Organoids

As with manual preparation of kidney organoids, iPSCs monolayers were differentiated for 7 days prior to being dissociated with Trypsin EDTA. The single cell suspension of differentiated cells was first counted using a Neubauer hemocytometer (BLAUBRAND catalog# BR7-18605) to obtain cell numbers prior to being centrifuged for 5 minutes at 200 x *g* to pellet cells in either a 50mL or 15mL polypropylene conical tube. This cell material was either transferred directly into a 100uL Gastight syringe (Hamilton Catalog# 7656-01) with a 21-gauge Removable Needle (Hamilton Catalog# 7804-12) for bioprinting, or resuspended to the working cell density with STEMdiff APEL or TESR-E6 media prior to transfer for bioprinting. In cases where cell slurry centrifugation was performed in the syringe, the prepared syringe was loaded into a proprietary adaptor to enable centrifugation at 400 x *g* within a 50mL polypropylene conical tube. Syringe/Adaptor assemblies were centrifuged for a total of 9 minutes to mirror the manual protocol. All syringes containing cellular bio-ink were loaded onto the NovoGen MMX bioprinter, primed to ensure cell material was flowing, and user-defined aliquots of bio-ink were deposited on to 0.4 µm polyester membranes of 6-well (Corning Costar catalog# 3450) or 96-well (Corning Costar catalog # 7369) Transwell permeable supports. Following bioprinting, organoids were maintained under the same conditions as those described above for manual organoid production.

### Histology

Kidney organoids were fixed overnight at 4°C in 2 or 4% paraformaldehyde (Electron Microscopy Sciences, Hatfield, PA). Fixed organoids were then pre-embedded in HistoGel (Thermo Fisher, Carlsbad, CA) and dehydrated and infiltrated with paraffin by automated processing with a TissueTek VIP tissue processing system (Sakura Finetek USA, Torrance, CA). Subsequent tissue blocks were cut in either a planar or transverse fashion to generate 5 micron thick, formalin-fixed, paraffin embedded (FFPE) sections on a Leica RM 2135 microtome (Leica Biosystems, Buffalo Grove, IL).

### H&E Staining

5 micron sections of FFPE kidney organoids were baked, deparaffinized, hydrated to water and stained following a standard regressive staining protocol using SelecTech staining solutions (Leica Biosystems, Richmond, IL; Haematoxylin #3801570, Define #3803590, Blue Buffer #3802915, and Eosin Y 515 #3801615). After staining was complete, the slides were serially dehydrated, cleared, and mounted in Permaslip (Alban Scientific Inc, St. Louis, MO #6530B). Histological images were acquired on a Zeiss Axio Imager A2 with Zeiss Zen software (Zeiss Microscopy, Thornwood, NY).

### FFPE Immunofluorescence

5 micron sections of FFPE kidney organoids were baked, deparaffinized, hydrated to water. Following rehydration, sections underwent heat-induced antigen retrieval in citrate buffer, pH 6.0 (Diagnostic BioSystems, Pleasonton, CA #KO35) and were blocked in 5% chick serum diluted in TBS-T (v/v). Tissues were incubated overnight with primary antibodies diluted in blocking buffer overnight. See Table 1 for antibody dilutions. All secondary antibodies were used at 1:200 (v:v).

### Whole Mount Immunofluorescence

Organoids were cut off Transwell filters and transferred to 48 well plates for fixation and immunofluorescence procedures. Fixation was performed using ice cold 2% paraformaldehyde (PFA; Sigma Aldrich) for 20 minutes followed by 15 minutes washing in three changes of phosphate-buffered saline (PBS). For immunofluorescence, blocking and antibody staining incubations were performed on a rocking platform for 3 hours at room temperature or overnight at 4°C, respectively. Blocking solution consisted of 10% donkey serum with 0.3% Triton-X-100 (TX-100; Sigma Aldrich) in PBS. Antibodies were diluted in 0.3% TX-100/PBS and details are supplied in Table 1. Primary antibodies were detected with Alexa Fluor-conjugated fluorescent secondary antibodies (Invitrogen), diluted 1:500. Organoids were washed in at least 3 changes of PBS for a minimum of 1 hour following primary and secondary antibody incubations. Imaging was performed in glass-bottomed dishes (MatTek) with glycerol-submersion using a Zeiss LSM 780 confocal microscope.

### Live imaging of reporter cell lines

Bioprinted organoids generated using the MAFB-mTagBFP2 iPS reporter cell line previously described ^17,18^ were live imaged for BFP2 intensity with an Apotome.2 fluorescent microscope (Carl Zeiss).

### Diameter Measurements

The cross-sectional diameter of the organoids was assessed over time by image-based analysis using ImageJ (version 1.51). Gross images were collected following print on Day 7 at a fixed distance with a 2x objective from plate surface. Each sample was manually outlined using the elliptical selection tool and used to calculate area in pixels for each image. Circular area values were converted to diameter in mm using the following equation:

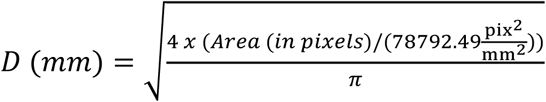

### Bioink Viability and Concentration Assay

Bioink viability and concentration were assessed by trypan blue exclusion and counted with the Nexcelom Cellometer. This was performed by sampling dispensed bioink before and after the printing of 2 rows (24 organoids) of a 96-well plate. Printed bioink was dispensed directly into 1.5mL Eppendorf tubes filled with APEL medium to dilute and counted. The Nexcelom results were placed into JMP for visualization and statistical analysis. A t-test was performed for analysis with only two conditions compared, a one-way ANOVA and Tukey comparison of means was performed for analysis with more than two conditions compared, and a bivariate fit was performed, using the fit mean, linear fit line, and 95% confidence interval to determine significant trends.

### Drug-Induced Nephrotoxicity Studies

Doxorubicin (Sigma-Aldrich, D1515) stock solution was prepared in DMSO and diluted into APEL media to the desired concentrations. Drug dosing was initiated after day 21 of the differentiation protocol. Dosing was performed by applying the full well volume of APEL medium ± doxorubicin to the apical basket of a Transwell permeable support (4mL for 6-well plates, 300 µL for 96-well plates) With media addition to the top, the organoids were fully submerged and exposed to any added compounds as the apical and basolateral compartments equilibrated. Drug-supplemented medium was replaced every other day until designated harvest time point.

## Organoid Viability Assessment

Kidney organoid viability following drug treatment was assessed by measuring ATP content with CellTiter-Glo or CellTiter-Glo 3D viability assays (Promega, Madison, WI, USA). In brief, harvested organoids from bioprinted in 6-well plates were individually loaded into Precellys tubes (Bertin Technologies, Bretonneux, France) with CellTiter-Glo buffer and dissociated using a Precellys 24 tissue homogenizer (Bertin Technologies, Bretonneux, France). Homogenized organoids were incubated at room temperature for 10 minutes, then centrifuged at 1000g for 2 minutes to separate buffer from homogenizing beads. Supernatants were transferred to a white opaque 96-well plate for luminescence measurement on a microplate reader (BMG Labtech, Germany). To analyze the ATP content in organoids bioprinted on 96-well plates, all media was aspirated and CellTiter-Glo 3D reagent was added to the apical chamber of Transwell permeable support. The plate was shaken at 400rpm for 5 minutes at room temperature, and then allowed to sit for 25 minutes prior to luminescent measurement in a white opaque 96-well plate on a microplate reader (BMG Labtech, Germany). Viability analysis was reported as percent of control by normalizing the ATP content of treated organoids relative to control organoids. Presented 6-well viability results are a composite of 3 studies with each normalized to respective control ATP levels within each study. Fitting of viability results was performed with GraphPad Prism 7.03 software (La Jolla, CA) using a four-parameter dose-response curve (Equation 1):

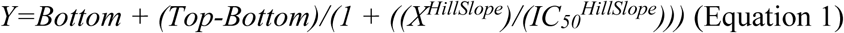

### Single cell transcriptional profiling and data analysis

Bioprinted and manual organoids were washed with PBS then dissociated by incubating in 500µl of 1µg/ml Liberase solution (Roche) on ice. Dissociation was aided by gentle pipetting with a p1000 tip at 5 minute intervals for a total of 20 minutes. Samples were then washed twice with DMEM containing 10% FCS to inactivate the Liberase enzyme and the cell clumps were removed by passing through 20µm sieve. Samples were centrifuged at 1300 rpm for 5 minutes and supernatant removed before resuspension in DMEM. Viability and cell number were assessed using automated cell counter (Thermofisher scientific) after trypan blue dye exclusion and samples were run in parallel on a Chromium Chip Kit (10x Genomics). Libraries were prepared using Chromium Single Cell Library kit V2 (10x Genomics), and sequenced on an Illumina HiSeq with 100bp paired-end reads.

The Cell Ranger pipeline (v1.3.1) was used to perform sample demultiplexing, barcode processing and single-cell gene counting ^25^. Data was imported in to Seurat (v2.3.1) for further quality control, clustering and marker analysis ^19,26^. Cells with less than 200 genes or greater than 6000 genes expressed, or greater than 15% mitochondrial gene content (defined by percent of cells with gene name beginning ‘MT’) were removed. Genes expressed in less than 3 cells were removed. Cell cycle prediction was performed for each cell using the cyclone function within the scran R package ^27,28^. Scaled data matrices were generated by regressing against the number of unique molecular identifiers (UMIs), percentage of mitochondrial genes expressed and cell cycle prediction scores for G1, S and G2M phase. Filtered datasets for standard organoids (1290 cells, mean 3435 genes detected) and bioprinted organoids (2381 cells, mean 3045 genes detected) were combined using canonical correlation analysis in Seurat. Clustering was performed at resolution 0.6 using 20 aligned canonical components calculated using a set of 11548 combined variable genes. Variable genes were defined for each dataset by average expression greater than 0.2 and less than 3 log normalised counts; dispersion greater than 1 and less than 10, with mitochondrial and ribosomal genes excluded. Differentially expressed marker genes of each cluster were identified in Seurat (Wilcoxon rank sum test) (Supplementary Tables 1 and 2).

Marker lists were exported and compared with published human single cell data ^29,30^, and established gene expression profiles from mouse ^31^. Gene ontology analysis was performed using TopFunn^32^.

## Figures

**Supplementary Figure 1.**
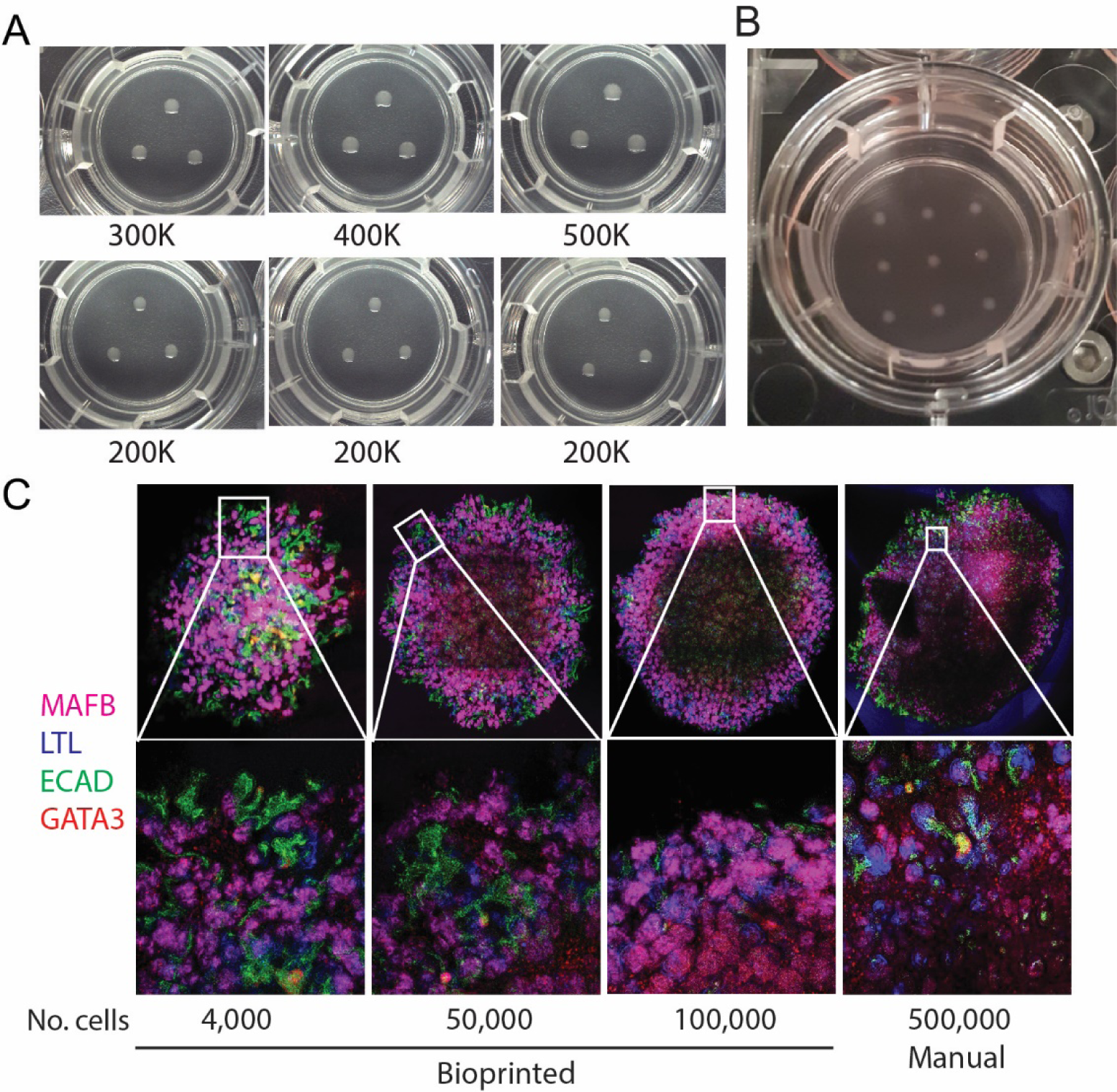
Bioprinted kidney organoids can be reliably and reproducibly generated from a variety of starting cell numbers. **A.** Transwell^®^ filters onto which triplicate kidney organoids have been bioprinted. The starting cell number is indicated. The top row illustrates a capacity to generate organoids with reducing numbers of cells. The bottom row illustrates the reproducibility of size when bioprinting a given cell number across multiple wells. **B.** 6 well Transwell with 9 bioprinted organoids each containing 96,000 cells. **C.** Immunofluorescence of bioprinted kidney organoids generated at total cell numbers as low as 4,000 cells per organoid showing the presence of podocytes (MAFB), proximal tubules (LTL) and distal tubule/collecting duct (ECAD, GATA3).

**Supplementary Figure 2.**
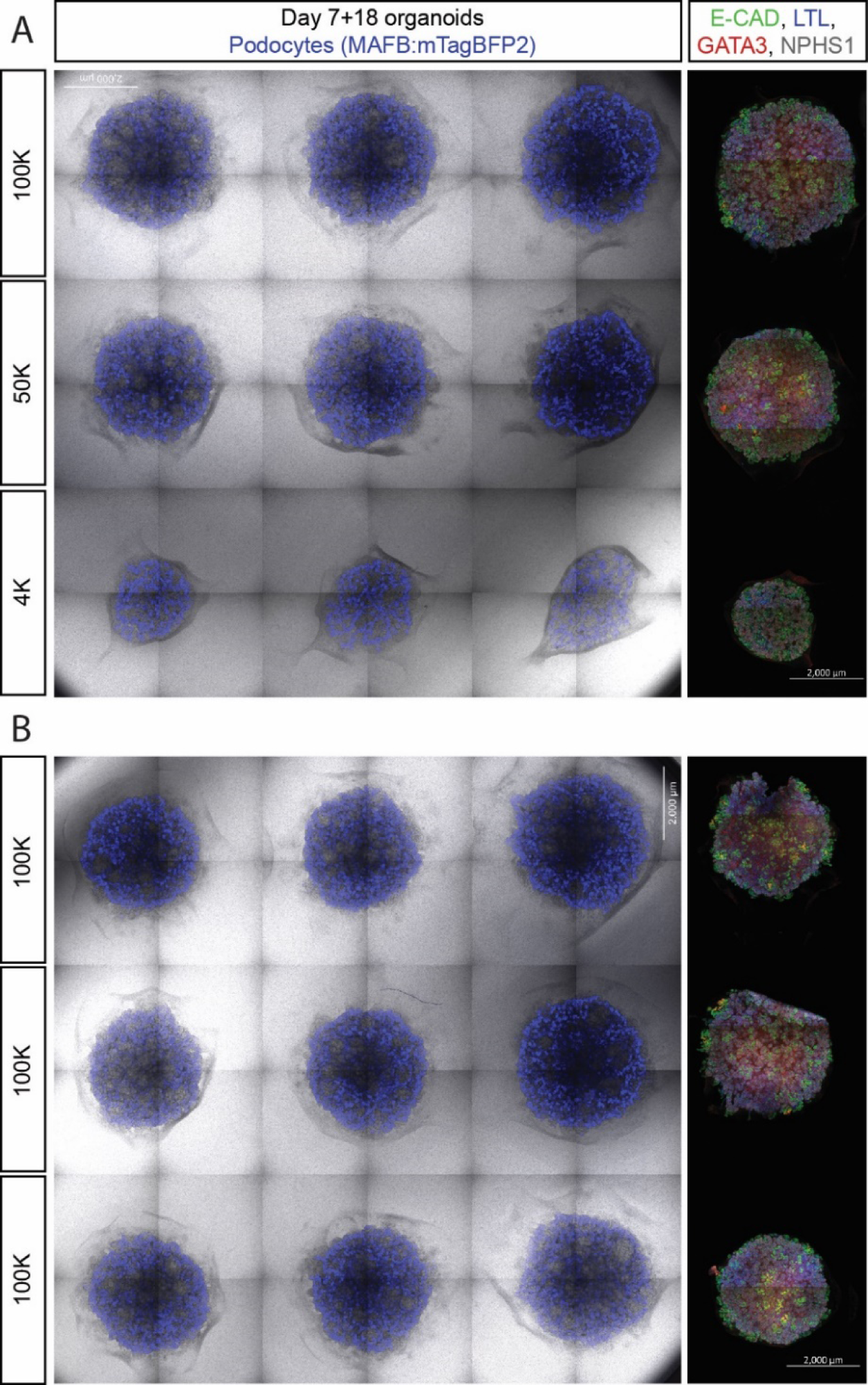
Kidney organoids bioprinted using a MAFBmTAGBFP reporter iPSC line showing podocyte differentiation (blue) with varying organoid size and differentiation reproducibility. **A.** Live phase contrast images showing endogenous MAFB fluorescence of organoids generated using cell numbers from 100K to 4K per organoid (left panel). Immunofluorescence analysis of day 7+18 organoids (right panel) reveals the consistent differentiation to podocyte (NPHS1, white), proximal tubule (LTL, blue), distal tubule (ECAD, green) and collecting duct (GATA3, red, and ECAD, green). **B.** Phase contrast/fluorescent images ofmultiple 100k bioprinted live MAFBmTAGBFP kidney organoids demonstrating differentiation reproducibility (left panel).. Immunofluorescence analyses (right panel) as for **B** above. Scale bars in **A** and **B** represent 2000µm.

**Supplementary Figure 3.**
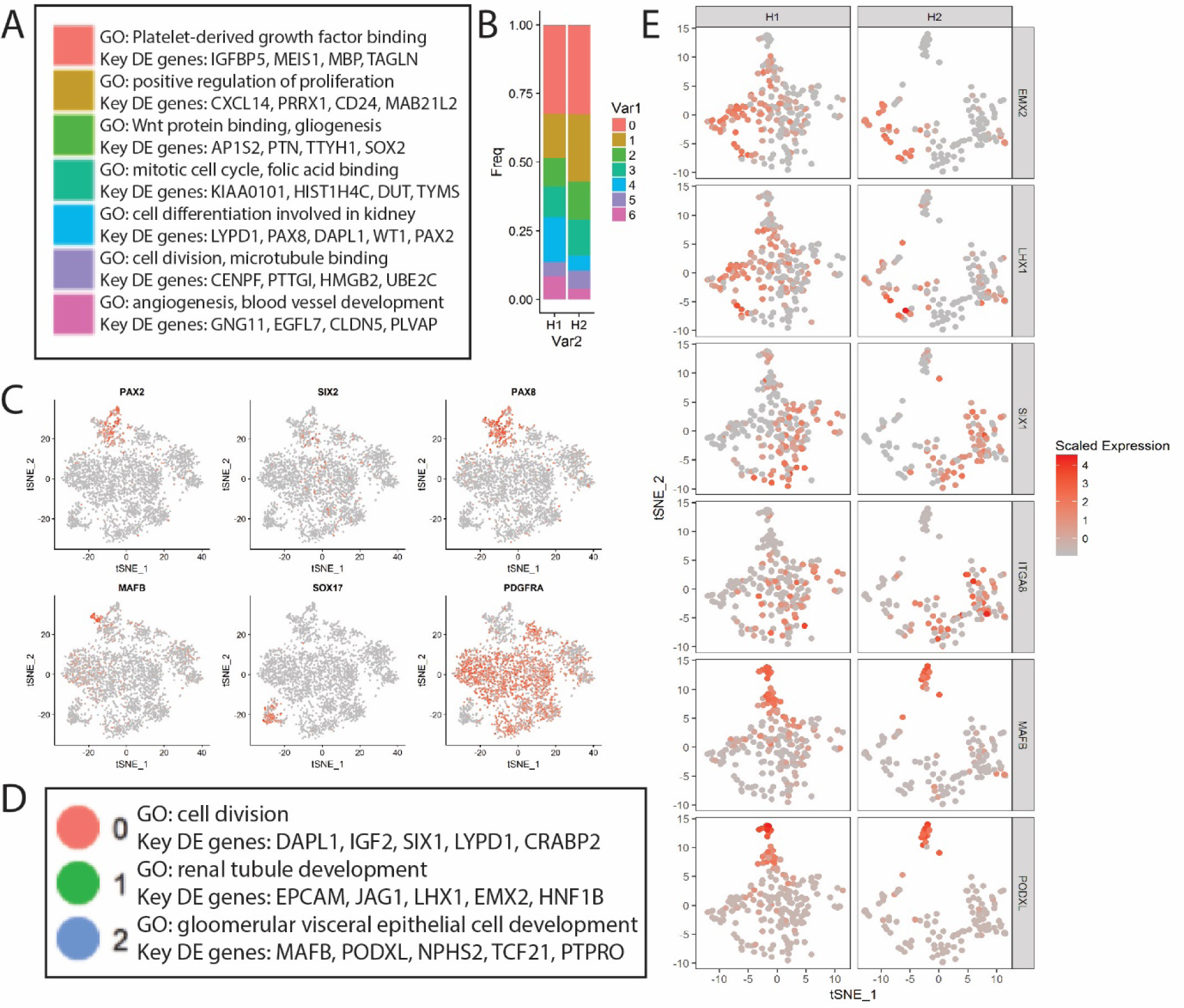
Single cell transcriptional profiling of standard and bioprinted organoids. **A.** GO terms and key differentially expressed genes for all clusters identified within single cell profiles of standard and bioprinted organoids. **B.** Relative cell numbers per cluster from standard versus bioprinted organoids. **C.** Feature plot showing the cells expressing key markers of nephron epithelium (PAX2, PAX8), nephron progenitors (PAX2, SIX1), podocytes (MAFB), endothelium (SOX17) and stroma (PDGFRA). **D.** GO terms and key differentially expressed genes for nephron subclusters 0, 1 and 2. **E.** Feature plot for key DE genes within nephron subclusters 0, 1 and 2 presented separately for the standard (H1, left) and bioprinted (H2, right) single cells.

**Supplementary Table 1. Differentially expressed (DE) genes for all clusters identified within standard and bioprinted organoids.**

**Supplementary Table 2. DE genes for all nephron subclusters identified within standard and bioprinted organoids.**

